# A new class of binding protein-dependent solute transporter exemplified by the TAXI-GltS system from *Bordetella pertussis*

**DOI:** 10.1101/2024.11.27.625746

**Authors:** Lily M. Jaques, Joseph F.S. Davies, Vanessa Leone, David J. Kelly, Christopher Mulligan

## Abstract

Tripartite ATP-dependent periplasmic (TRAP) transporters are widespread in prokaryotes, but absent in eukaryotes, and facilitate the uptake of a wide variety of substrates. TRAP transporters are composed of a substrate binding protein (SBP) and two unequally sized membrane components that exist as either separate proteins or are fused into a single polypeptide. Almost all TRAP SBPs exist as monomeric globular proteins that bind substrate and present it to the membrane component. Here, we describe the discovery and characterisation of a novel TRAP SBP from the TAXI subfamily with a previously unidentified architecture. BP0403 from the human pathogen *Bordetella pertussis* is a predicted lipoprotein composed of 3 distinct domains; an α/β globular domain with a unique fold, a long helical domain and a C-terminal TAXI SBP domain. Expression and purification of the full-length protein reveals that it forms a stable dimer. Structural modelling of the dimer interface and molecular weight analysis using size exclusion chromatography of the individual domains reveals that the interdomain helical region is solely responsible for dimerization. Differential scanning fluorimetry (DSF) and intrinsic tyrosine fluorescence reveal that BP0403 binds L-glutamate with nanomolar affinity. Unexpectedly, analysis of the genome context of BP0403 reveals the complete absence of characteristic genes for TRAP membrane components but co-localisation and translational coupling with *gltS*, encoding a Na^+^/glutamate symporter. In other bacteria, we identified fused BP0403-GltS homologues, strongly suggesting that this constitutes a completely novel SBP-dependent secondary active transporter. Structural comparisons suggest GltS operates by an elevator-type mechanism, like TRAP transporters; the association of an SBP with this class of secondary transporter is an emerging theme.

## Introduction

Substrate binding proteins (SBPs) are obligate components of many solute transporters in prokaryotes, where they deliver solutes to integral membrane transporters, often conferring stringent selectivity, high affinity and directionality to solute uptake. SBPs have been recruited by three main transporter families; the primary-active ATP-binding cassette (ABC) transporters that hydrolyse ATP to power transport^1^, and the secondary-active tripartite tricarboxylate transporter (TTT) and tripartite ATP-independent periplasmic (TRAP) transporter families, both of which harness electrochemical ion-gradients to drive transport^2–4^.

TRAP transporters (TCDB ID: 2.A.56) are the largest and best characterised SBP-dependent secondary active transporter family and contribute to virulence of several human pathogens^4^. TRAP transporters are composed of an SBP and two unequally sized membrane components that can exist as two separate proteins or fused into a single polypeptide^2,5^. The TRAP family can be further divided into two subfamilies; the DctP-type and the TAXI (<u>T</u>RAP <u>a</u>ssociated e<u>x</u>tracytoplasmic immunogenic) TRAP transporters; both subfamilies have homologous membrane components, but the SBPs share no amino acid sequence similarity^2^. DctP-type TRAPs were the first discovered and are named after the first TRAP transporter identified, DctPQM from *Rhodobacter capsulatus*, which is C4-dicarboxylate specific^5^. DctP-type TRAP systems have been extensively characterised, revealing a diverse portfolio of substrates, most commonly aliphatic or aromatic organic acids^6–9^.

Recent structural characterisation of fused DctP-type TRAP membrane components reveal that they are monomeric proteins composed of two domains, a scaffold domain and a transport domain, the latter of which houses all substrate binding residues^10–12^. TRAP membrane components share the same fold as members of the divalent anion Na^+^ symporter (DASS) family^13–15^. As such, they also share a general transport mechanism in which the transport domain undergoes an elevator-like movement through the membrane to facilitate transport^10–12,16,17^. Similarly to DASS family members^18^, liposome reconstitution of DctP-type TRAP systems has revealed that transport is dependent on the presence of a Na^+^ gradient^11,12,19,20^.

The TAXI TRAP subfamily is relatively poorly characterised in comparison, with only a handful of substrates known or postulated and only two experimentally determined structures currently published^21,22^. The TAXI TRAP systems differ in several ways from their DctP-type cousins. Whereas DctP-type TRAP transporters can have either fused or separate membrane components, TAXI systems invariably have a fused membrane protein^2^, and while both DctP-type and TAXI TRAP systems are widespread in bacteria, the TAXI group are the *only* TRAP systems found in archaea. In contrast to the Na^+^-dependence of DctP-type transporters, at least one TAXI TRAP transporter harnesses the proton-motive force to drive transport^23^. In addition to these differences, a higher proportion of TAXI SBPs appear to be lipoproteins compared to DctP-type TRAP SBPs, although the significance of this is not clear^23^.

Although DctP-type and TAXI SBPs are unrelated at the sequence level their domain architecture and overall structures are similar. They are usually monomeric, 300-350 amino acids long and are invariably solely composed of 2 α/β globular domains and a hinge region, which closes upon interaction with substrate in a “venus flytrap”-like mechanism^24,25^. While there are examples of ‘orphan’ TRAP SBPs, the vast majority of the genes encoding them are operonic, with the genes encoding the characteristic TRAP membrane proteins with which they interact. However, in this work we describe the discovery and characterisation of a novel elongated, dimeric TAXI SBP from the human pathogen *Bordetella pertussis*, which has recruited an additional globular domain. We present compelling evidence that this unusual TAXI SBP binds glutamate but is not part of a TRAP transporter; rather, it utilises a member of the unrelated GltS family of Na^+^ coupled glutamate symporters for uptake. This conclusion is strengthened by the identification of TAXI-GltS fusion proteins encoded in the genomes of other bacteria. This unprecedented architecture reveals for the first time the existence of a novel, previously uncharacterised type of SBP-dependent transporter.

## Methods

### Molecular Biology

Genes encoding full-length BP0403 (BP0403_FL_), BP0403 N-terminal domain (BP0403_NTD_) and BP0403 C-terminal domain (BP0403_SBP_) were codon optimised, synthesised and cloned into pET151 by Invitrogen GeneArt. For both BP0403_FL_ and BP0403_NTD_, the N-terminal signal peptide (MTMFIRWLILSACLLLAAC) was not included in the final construct.

### Protein expression and purification

To express BP0403_FL_, BP0403_NTD_ and BP0403_SBP_, *E. coli* BL21 (DE3) was transformed with a pET151 plasmid with the gene encoding these proteins in frame with an N-terminal His_6_ affinity purification tag. The expression strains were grown in LB media supplemented with 100 µg/ml ampicillin in a 2.5 L baffled Tunair flask. The cultures were grown at 37°C to an OD600 of 0.8, at which point expression was induced by addition of 1 mM IPTG. Cultures were incubated for a further two hours, then harvested by centrifugation at 4000 rcf for 20 min and resuspended in Lysis buffer (50 mM Tris-HCl, pH 8, 200 mM NaCl, 5% glycerol). Cells were lysed by ultrasonication and the lysate was clarified by centrifugation at 20 000 rcf for 20 min at 4°C.

The clarified lysate was applied to Ni-NTA resin (Qiagen) at room temperature. The resin was washed with 20 column volumes (CV) of Wash Buffer (50 mM Tris-HCl, pH 8, 100 mM NaCl, 5% glycerol, 20 mM imidazole), and the bound protein was eluted with 5 x 0.5 CV of Elution Buffer (50 mM Tris-HCl, pH 8, 100 mM NaCl, 5% glycerol, 300 mM imidazole). The protein was further purified using size exclusion chromatography with a Superdex 200 Increase 10/300 GL column (GE Healthcare) equilibrated with SEC buffer (50 mM Tris-HCl, pH 8, 100 mM NaCl, 5% glycerol).

### SEC-based oligomeric state analysis

For oligomeric state analysis, IMAC purification elution fractions were pooled and concentrated, incubated at room temperature for 20 min with 5 mM EDTA and 100 mM glycerol, and applied to a Superdex 200 Increase 10/300 GL column equilibrated with SEC buffer. SEC was performed at a flow rate of 0.5 ml/min with SEC buffer. The calibration curve was generated by determining the elution volumes of SEC standards (Thermo Scientific) applied to the same column under the same conditions and plotting the elution volumes as a function of log molecular weight of the standards.

### Differential Scanning Fluorimetry (DSF)

DSF was performed by mixing the protein of interest, either 1.5 µM BP0403_FL_, 19 µM BP0403_SBP_, or 6.5 µM BP0403_NTD_, with 1x SYPRO Orange dye (Merck) in DSF buffer (50 mM Tris-HCl, pH 8, 20 mM NaCl). Potential ligands were added to a final concentration of 1 mM, where appropriate. A QuantStudio 3 RT-PCR thermocycler (Invitrogen) was used to initially cool to 5°C at a rate of 1.6 °C/s. The temperature was then increased to 95 °C at 1.6 °C/s and readings were taken every 1°C while holding the temperature for 5 s using the SYBR reporter dye setting. DSF data was exported to Microsoft Excel and GraphPad Prism for analysis and presentation.

### Intrinsic tyrosine fluorescence

Fluorescence assays was measured using a Cary Eclipse Fluorimeter (Aglient) in a 2 ml quartz cuvette. Ligand titrations were performed using 50 nM SEC-purified BP0403_SBP_ in time-based acquisition mode with excitation and emission wavelengths of 280 and 310 nm, respectively. Excitation slit width was set to 5 nm, emission slit was set to 10 nm, the photomultiplier tube (PMT) voltage was set to 950 V. Emission spectra were collected using 500 nM SEC-purified BP0403_SBP_ with an excitation wavelength of 280 nm, and an emission range of 280-400 nm. Excitation slit width was set to 5 nm, emission slit was set to 10 nm, the photomultiplier tube (PMT) voltage was set to 800 V, and a Savitzky-Golay smoothing factor of 15 was applied.

### Protein modelling

Structural models of BP0403 and BPP3828 monomers were extracted from the AlphaFold Protein Structure Database^26^. The BP0403 dimer and GltS dimer were modelled using ColabFold^27^. Structural models of the BP0403:GltS heterotetramer were built using AlphaFold-Multimer v2.3.0^26,28^ (Evans et al., 2022; Jumper et al. 2021). The sequences of BP0403 (accession ID: Q7VSK8) and GltS (accession ID: Q7VSK9) of *B. pertussis* were taken from the UniProt database. Five predictions were obtained, one for each neural network model, and we selected the one with the highest combined predicted Template Model (pTM) and interface pTM score. All images were generated using UCSF ChimeraX^29^.

## Results

### Elongated TAXI SBPs are predicted to have multiple domains

In our previous analysis of the distribution and variation of TAXI SBPs^22^, we observed that one SBP, BP0403 from *Bordetella pertussis*, was substantially larger than the other TAXI proteins in our sequence alignment. While all other TAXIs SBPs in the alignment were 300-350 amino acids long, similar to DctP-type SBPs^2^, BP0403 was composed of 503 amino acids. Intrigued by this surprisingly long TAXI SBP, we investigated its predicted structure using the AlphaFold2 server^26^.

The BP0403 model structure revealed a large globular protein that could be divided into 3 distinct domains (Fig. 1A); an N-terminal α/β globular domain (NTD), a long helical domain (HD), and a large C-terminal α/β globular domain composed of a standard SBP architecture (SBP domain). To confirm that the C-terminal domain indeed has an SBP fold, we aligned the predicted structure of the BP0403 SBP domain with the known structure of the glutamate-specific TAXI SBP, VcGluP^22^, which superimposed with an RMSD of 3.4 Å across 293 residues, confirming its identity (Fig. 1B). As all other SBPs are secreted into the periplasm or are lipid-anchored periplasmic-facing proteins^2^, we analysed the BP0403 amino acid sequence using SignalP 6.0^30^, which revealed the presence of an N-terminal lipoprotein signal peptide (Sec/SPII) that would tether the protein to the outer leaflet of the inner membrane (Fig. 1A).

**Figure 1.**
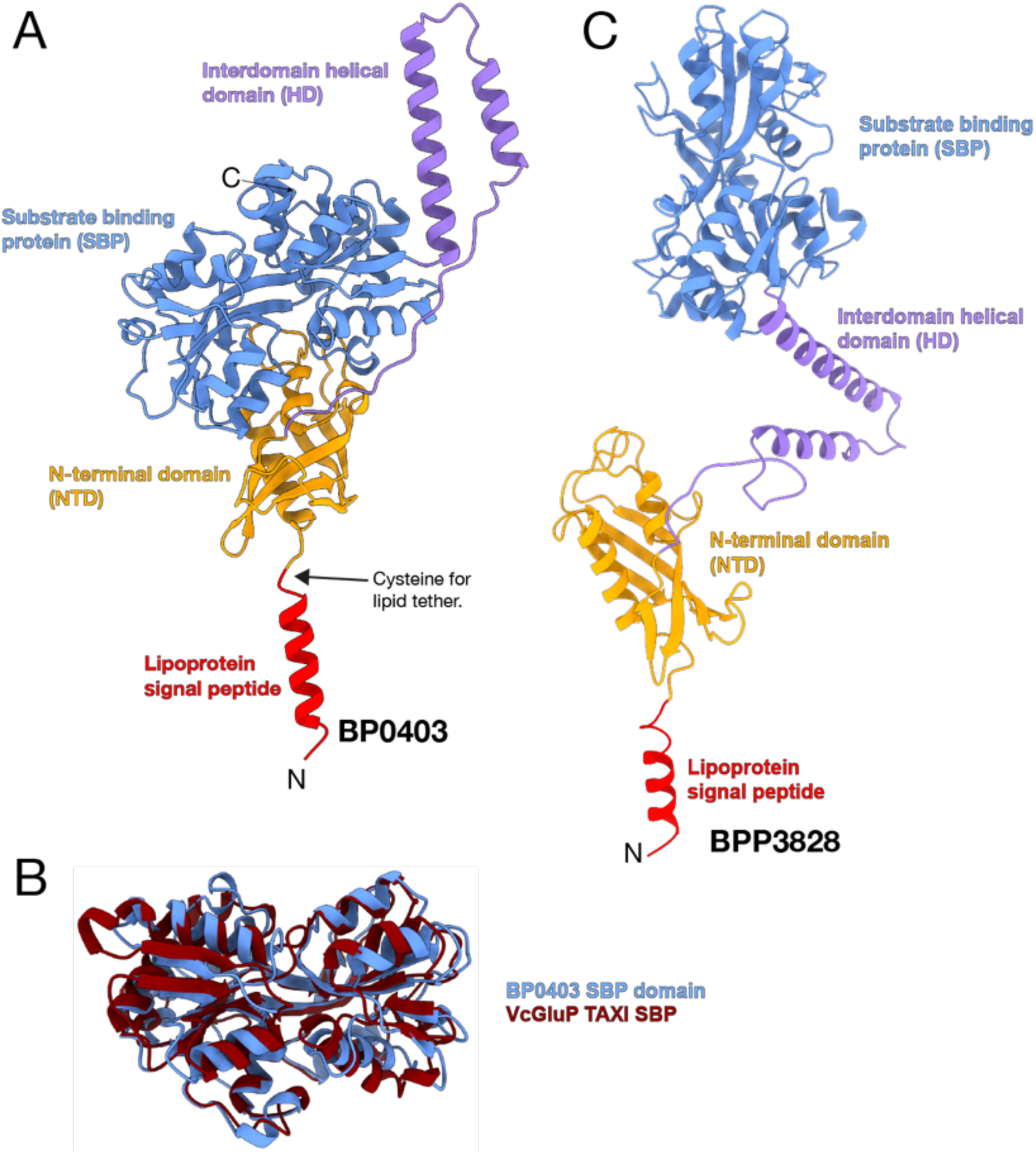
Structural model of BP0403. **A)** AlphaFold2 model of BP0403 colour coded as follows; lipoprotein signal peptide (red), N-terminal domain (orange), interdomain helical region (purple), and SBP domain (blue, LDDT representation is in SI Fig. 1A) **B)** Superimposition of the SBP domain from BP0403 and the crystal structure of glutamate-specific TAXI, VcGluP (PDB ID: 8S4J). **C)** AlphaFold2 structure of BPP3828 from *B. parapertussis* revealing an elongated structure (same colour code as in **A**).

To shed light on the potential domain arrangement in BP0403, we analysed the AlphaFold2 model of the BP0403 homologue BPP3828 from *Bordetella parapertussis*, which modelled the helical domain in a more extended arrangement (Fig. 1C). This arrangement suggests a possible mechanism in which the SBP is tethered to the membrane but has substantial freedom of movement due to the flexible helical domain.

### BP0403 binds L-glutamate

To investigate the structural and functional characteristics of BP0403, we expressed the gene in *E. coli* with an N-terminal histidine tag in place of the lipoprotein signal peptide. Overexpressed BP0403 was purified using IMAC, which revealed an intense band on SDS-PAGE at ∼50-55 kDa, consistent with the molecular weight of 53.7 kDa estimated from the amino acid sequence (Fig. 2A, inset). Further purification using size exclusion chromatography (SEC) revealed a single symmetrical peak, indicative of folded and stable protein (Fig. 2A).

**Figure 2.**
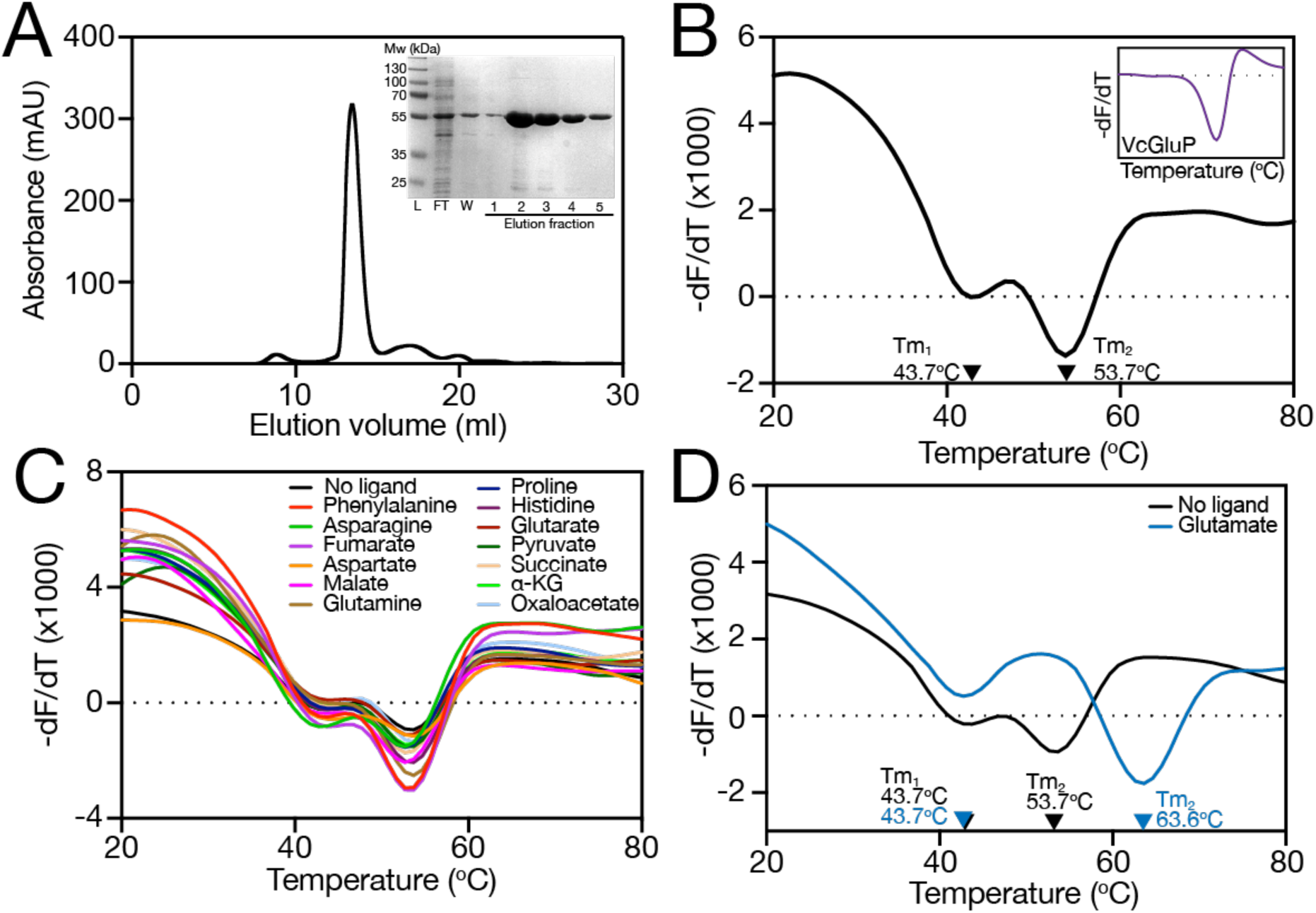
Purification and DSF analysis of BP0403. **A)** Size exclusion chromatography (SEC) trace of IMAC purified BP0403. *Inset*: SDS-PAGE of IMAC fractions from BP0403 purification, including molecular weight ladder (L), flowthrough (FT), the wash fraction (W), and elution fractions 1-5. **B-D)** Derivatives of the unfolding curves (dF/dT) for **B)** BP0403 in the absence of ligand and VcGluP in the absence of ligand (inset), **C)** for BP0403 in the presence of several potential ligands, and D) in the presence and absence of L-glutamate. Derivative curves shown are representative curves taken from quadruplicate datasets. Each assay was performed on at least 2 occasions with the same outcome.

To investigate the ligand specificity of BP0403 we used differential scanning fluorimetry (DSF), which provides a readout of the melting temperature (Tm) of the protein, which often increases upon ligand binding, due to increased protein stabilisation^31^. DSF analysis of conventional TAXI SBPs consistently produces a single trough, indicative of a single melting event, for example, with the L-glutamate specific TAXI SBP, VcGluP (Fig. 2B, *inset*)^22^. However, analysis of BP0403_FL_ using DSF revealed an unusual double dip pattern (Fig. 2B), suggesting the occurrence of two melting events with separable Tms of 43.7°C (Tm_1_) and 53.7°C (Tm_2_), which we reasoned likely reflects the independent melting of the globular SBP and N-terminal domains.

To identify the ligand binding specificity of BP0403_FL_, we screened a library of 14 different compounds consisting of a range of organic anions, with similar features to “classical” TRAP ligands, using DSF (Fig. 2C and D). While the majority of the compounds induced no change in the position of either of the melt troughs of BP0403_FL_ (Fig. 2C), we observed a pronounced rightward shift of only Tm_2_ in the presence of L-glutamate (Fig. 2D). These data strongly support the conclusion that Tm_2_ relates to the SBP domain and identifies L-glutamate as the ligand.

To determine which melt trough belonged to which globular domain, we performed DSF on the isolated domains, BP0403_NTD_ (residues 20-139) and BP0403_SBP_ (residues 208-503). DSF analysis revealed that each isolated domain produces a single melt trough; BP0403_SBP_ with a Tm of 61.6°C, and BP0403_NTD_ with a Tm of 39.8°C (Fig. 3A). While neither of the Tms from the isolated domains corresponds exactly with the two Tms obtained from BP0403_FL_, comparison of these data with the double dip DSF data of BP0403_FL_ indicates that Tm_1_ likely corresponds with the BP0403_NTD_, and Tm_2_ is likely the BP0403_SBP_ (Fig. 3A).

**Figure 3.**
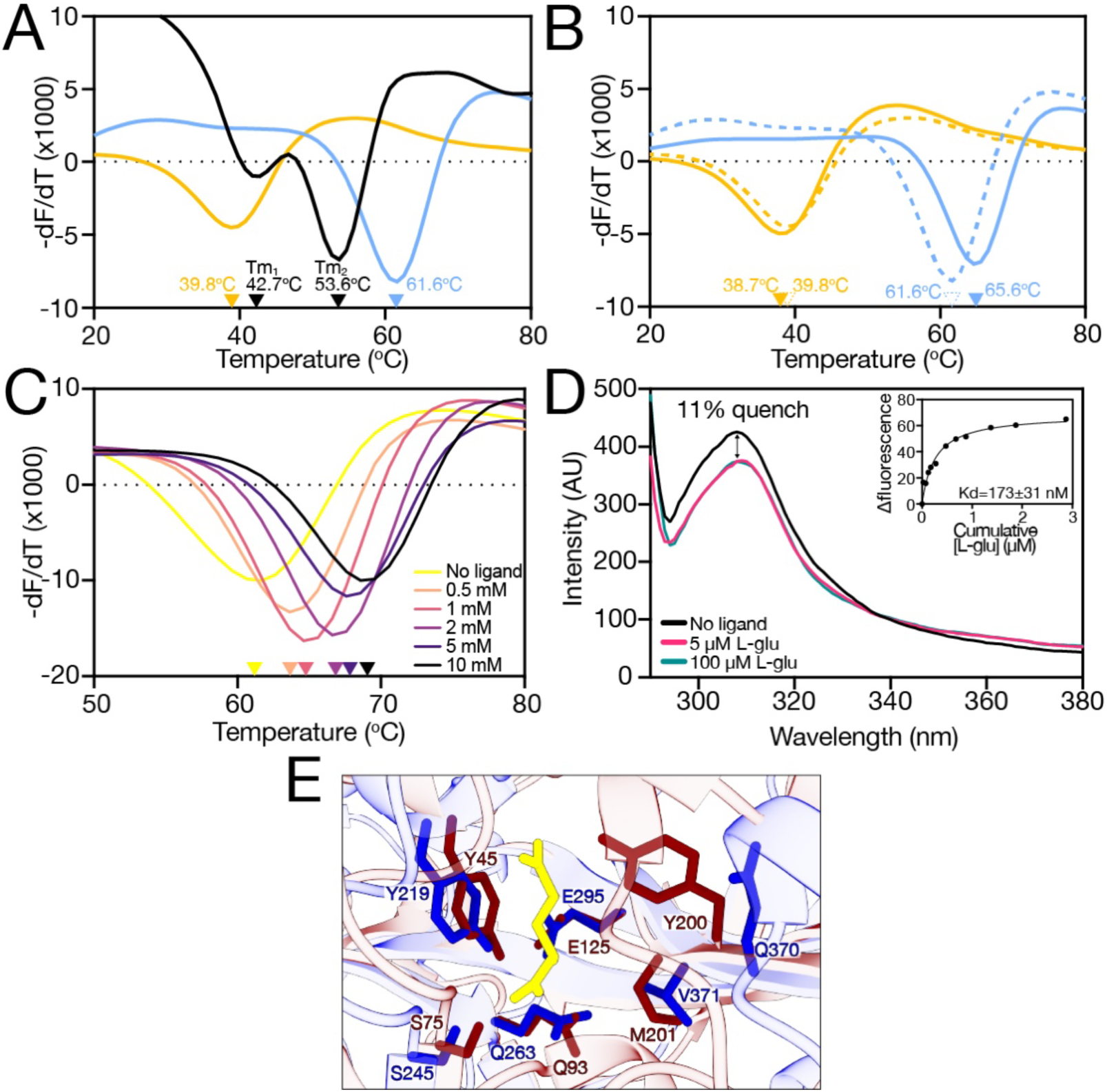
Ligand binding analysis on isolated SBP and NTD domains. **A)** Derivatives of the unfolding curves (dF/dT) for BP0403_FL_ (black data), BP0403_SBP_ (blue data) and BP0403_NTD_ (orange data). **B)** Derivatives of the unfolding curves (dF/dT) for BP0403_SBP_ and BP0403_NTD_ with (solid lines) and without (dashed lines) 1 mM L-glutamate. **C)** Derivatives of the unfolding curves (dF/dT) for BP0403_SBP_ with increasing concentrations of L-glutamate. **D)** Tyrosine fluorescence scan of BP0403_SBP_ in the absence of ligand (black data), 5 µM L-glutamate (pink data), 100 µM L-glutamate (cyan data). *Inset*: representative L-glutamate binding curve. Average Kd is shown (n=11) and error represents SEM. **E)** Superimposition of the binding site residues of BP0403 AF2 model (blue residues) and the VcGluP crystal structure (red residues).

To confirm that Tm_2_ corresponds with the BP0403_SBP_, we performed DSF individually on the BP0403_NTD_ and BP0403_SBP_ in the presence L-glutamate (Fig. 3B). While addition of 1 mM L-glu had essentially no effect on the Tm of BP0403_NTD_, the Tm of the BP0403_SBP_ was increased by 4°C (Fig, 3B), substantially less than the 9.9°C shift seen for BP0403_FL_’s Tm_2_ (Fig. 2D). To investigate this further, we performed an L-glutamate titration with BP0403_SBP_ using DSF, which revealed the dose-dependent Tm increase expected for ligand binding to an SBP (Fig. 3C). To support the DSF-based ligand identification, we wished to perform tryptophan fluorescence spectroscopy on BP0403_FL_. However, while BP0403_FL_ contains 2 tryptophans, they are both located in the NTD, which does not bind ligand according to our DSF data. Specifically monitoring tyrosine fluorescence with BP0403_FL_ was not possible due to the tryptophan emission masking any dose dependent fluorescence changes. Therefore, we monitored tyrosine fluorescence changes with the isolated SBP domain (Fig. 3D). Addition of 5 µM L-glutamate induced an 11% quench in the protein fluorescence, which was saturable, as there was no further quenching upon addition of 100 µM L-glutamate (Fig. 3D). We performed an L-glutamate titration that revealed a K_d_ of 180 ± 32 nM for L-glutamate (Fig. 3D, inset).

To investigate the amino-acid binding determinants of BP0403_SBP_, we superimposed the model of BP0403_SBP_ with the X-ray structure of VcGluP, a TAXI SBP of conventional size from *Vibrio cholerae* that we previously showed stereoselectively binds L-glutamate with high-affinity and L-glutamine and pyroglutamate with lower affinity^22^. This structural comparison revealed an almost identical arrangement of several amino acids in the predicted substrate binding site (Fig. 3E), including Y219 and Q263, which are equivalent to the two most influential binding site residues in VcGluP (Y45 and Q93 in VcGluP)^22^. This similarity of binding site architecture further supports the ligand specificity of BP0403 we have identified.

### BP0403 is a dimer

Comparison of our DSF analysis of BP0403_FL_ and BP0403_SBP_ revealed that 1) the isolated SBP domain has a substantially higher Tm (∼8°C) than when it is part of the full-length protein, and 2) the ligand-induced ΔTm is much smaller for the isolated SBP domain than when part of BP0403_FL_ (4°C vs 9.9°C, respectively). These data suggest that there is a fundamental difference in the arrangement of the SBP domain when isolated compared to when it is part of the full-length protein, and contrary to our working model (Fig. 1), the DSF data do not support a tethered, but a freely mobile SBP. We reasoned that this could be due to oligomerisation of BP0403, which is a very rare, but not unheard-of arrangement for TRAP SBPs^32^.

To investigate the oligomeric state of BP0403, we applied purified BP0403_FL_ to a SEC column and compared its elution volume to the elution volumes of protein standards of known molecular weight (Fig. 4A). Analysis of the molecular weight of BP0403 using a standard curve generated from the molecular weight standards revealed that the elution volume corresponds to a 114.8 kDa protein (Fig. 4B). As the estimated molecular weight of BP0403 is 53.7 kDa based on the amino acid sequence, these data are consistent with BP0403 forming a stable dimer in solution.

**Figure 4.**
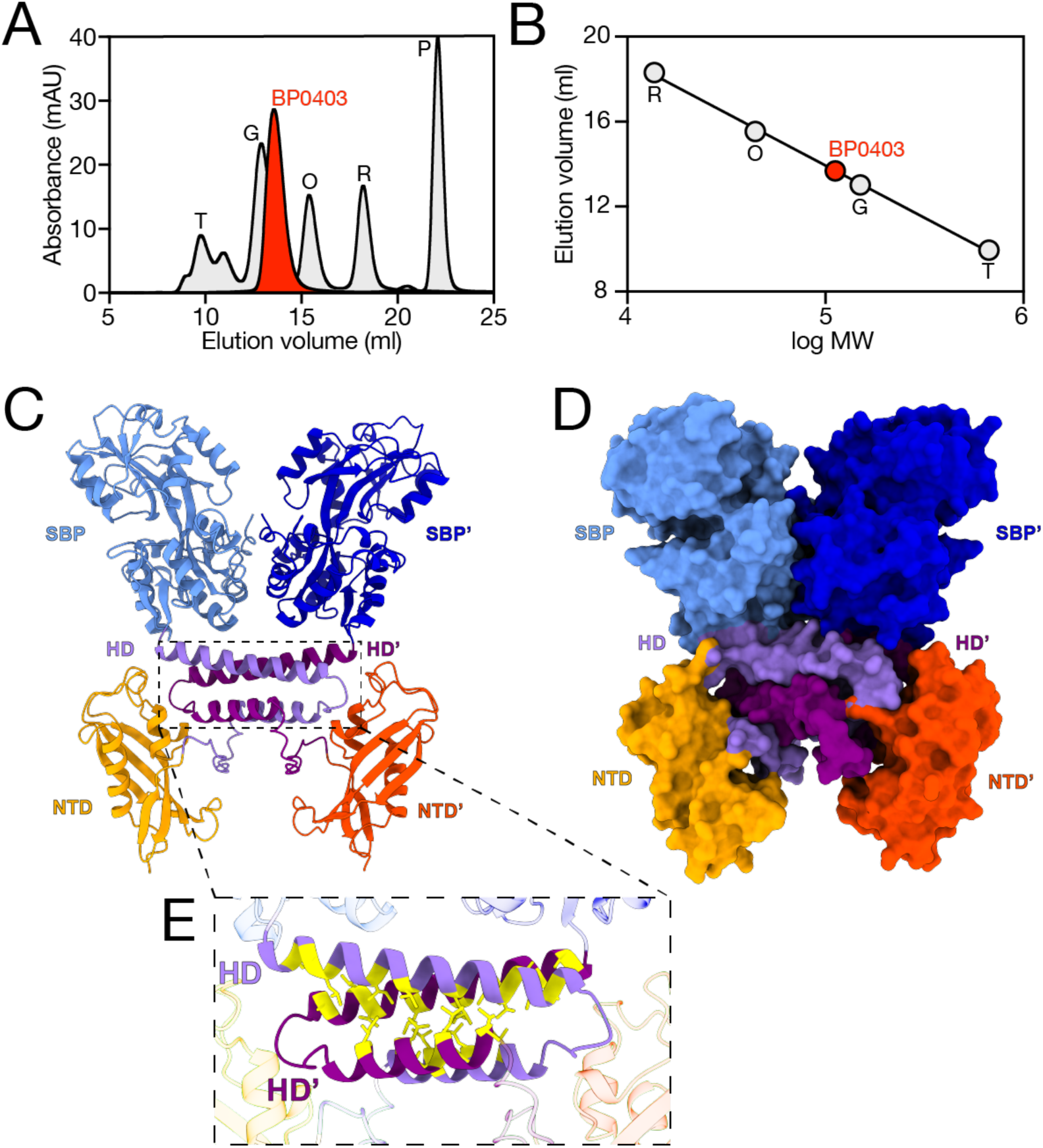
Oligomeric state analysis of BP0403. **A)** SEC trace of BP0403 (red data) compared to 5 SEC standards of known molecular weight (grey data); thyroglobulin (T, 660 kDa), gamma-globulin (G, 150 kDa), ovalbumin (O, 43 kDa), ribonuclease A (R, 14 kDa) and p-aminobenzoic acid (P, 0.14 kDa). **B)** Elution volume as a function of log MW of the SEC MW standards (grey) and BP0403 (red). **C)** Cartoon representation of the Colabfold model of a BP0403 dimer (LDDT representation is in SI Fig. 1B). Domains are coloured as in Figure 1; protomer 1 is lighter colours compared to protomer 2. **D)** Surface representation of BP0403 dimer model. **E)** Close-up image of the helical domain interactions. Hydrophobic residues are highlighted in yellow. For C) and D) the signal peptide has been omitted from analysis.

Intrigued by this finding, we modelled the BP0403 dimer using Colabfold^27^, which predicted minimal interfacial contacts between the SBP domains in the dimer and no contacts between the NTDs (Fig. 4C and D). Indeed, the majority of interprotomer contacts come from a complex arrangement of the helical domains, which are predicted to form interlocked hairpins (Fig. 4C-E). The 4 helices contributing to this interface are amphipathic and the majority of the interfacial interactions are hydrophobic (Fig. 4E).

As our dimer model predicts that the helical domains facilitate all the interfacial contacts, it follows that the isolated SBP and NTD domains should be monomeric. To test this prediction, we generated truncations of BP0403 and independently expressed either BP0403_SBP_ or BP0403_NTD_.

We purified each of the independent domains and SDS-PAGE analysis revealed that each domain migrated in-line with its estimated molecular weight based on the amino acid sequences; 33.8 kDa and 16.4 kDa for BP0403_SBP_ and BP0403_NTD_, respectively (Fig. 5A and B). We further purified each domain using SEC and compared the elution volumes to those of the molecular weight standards used previously (Fig. 5C and D). This analysis revealed single peaks for BP0403_SBP_ and BP0403_NTD_ that correspond with proteins of 34.5 kDa and 18.9 kDa, respectively (Fig. 5C and D). These data demonstrate that the isolated domains are monomeric in solution and fully support a model in which the dimer interface is formed via interaction between the helical domains.

**Figure 5.**
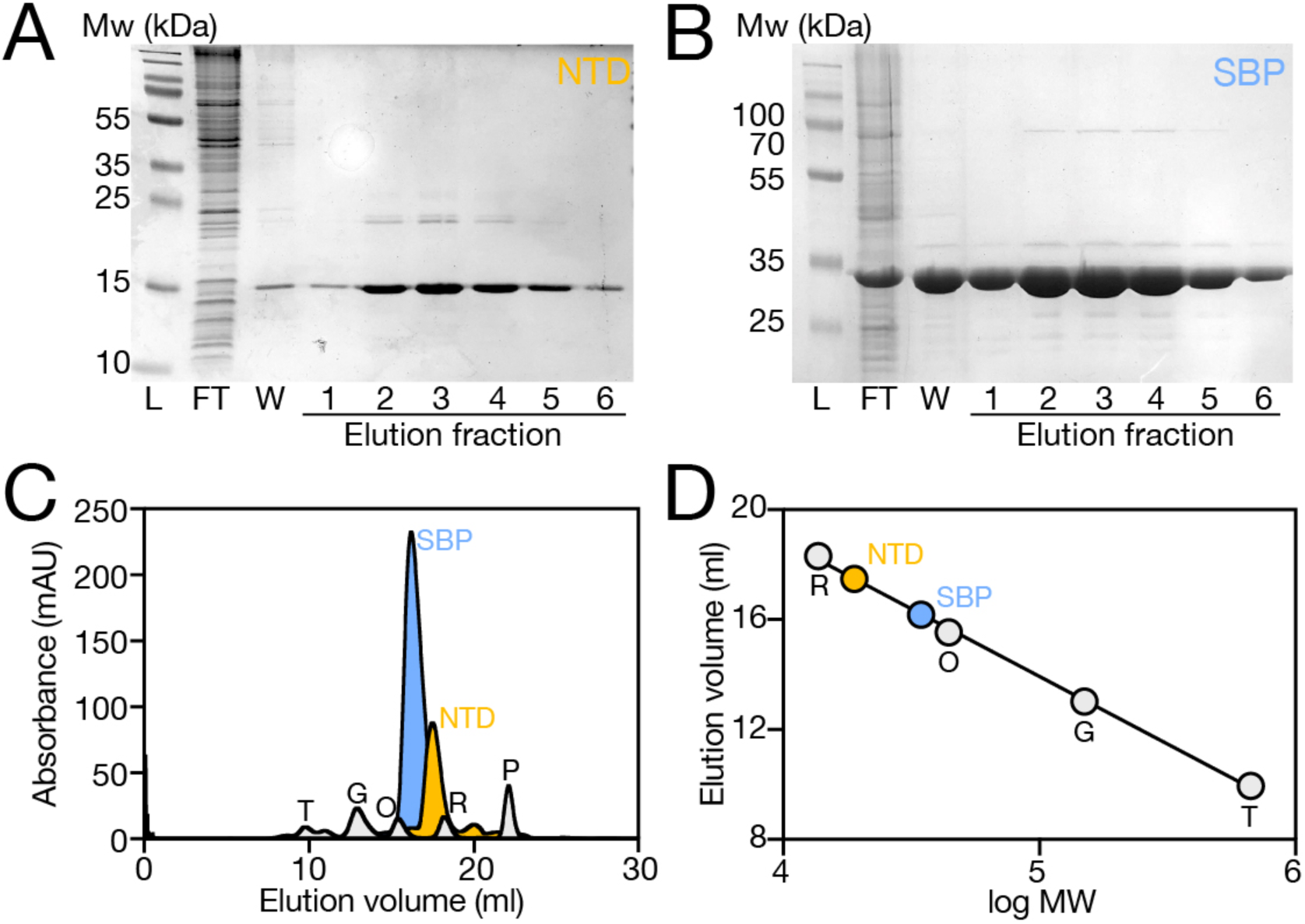
Oligomeric state analysis of isolated SBP and NTD domains. **A) and B)** SDS-PAGE analysis of the IMAC purification fraction for BP0403_NTD_ and BP0403_SBP_, respectively. Lane labels are the same as Figure 2A. **C)** SEC trace of BP0403_SBP_ (blue data) and BP0403_NTD_ (orange data) compared to 5 SEC standards of known molecular weight (grey data); same standards as in Figure 4. **D)** Elution volume as a function of log MW of the SEC MW standards (grey data), BP0403_SBP_ (blue data), and BP0403_NTD_ (orange data).

### BP0403 is not associated with a TRAP membrane component but with GltS

TRAP SBPs must interact with an integral membrane transporter to move substrates across the cytoplasmic membrane. While some TRAP SBP genes are ‘orphans’^2^, the vast majority are co-localised in an operon with the gene(s) expressing the TRAP membrane component(s). Therefore, to identify the cognate membrane component(s) to accompany BP0403, we analysed the genome context of *BP0403*. To our surprise, there were no genes encoding TRAP membrane proteins in the *BP0403*-containing operon or immediate gene region. Instead, *BP0403* is downstream of *BP0402*, which encodes *gltS*, with the end of *gltS* overlapping with the start of *BP0403* by 3 bp, indicative of translational coupling (Fig. 6A). GltS belongs to the glutamate:Na^+^ symporter (ESS) family (TCDB ID: 2.A.27^33^), members of which are widespread in prokaryotes, but are not found in eukaryotes. Further analysis of genome context data using SeedViewer revealed a gene encoding GltS was also adjacent to genes encoding the elongated TAXI SBPs from multiple organisms (Fig. 6A). Furthermore, extending our search to include TAXI SBPs of conventional size, we found several operons in which a TAXI SBP gene was directly up- or downstream of a gene encoding GltS (Fig. 6B). Importantly, this genetic association is found in multiple disparate organisms across the prokaryotic phylogeny, strongly suggesting that the TAXI SBP and GltS are functionally coupled. Low resolution cryo-EM and blue native PAGE of *E. coli* GltS revealed that it is a dimer and functional characterisation showed preferential transport of L-glutamate, with D-glutamate and 2-methyl glutamate transported at much lower affinity^34,35^, strengthening the link with our dimeric L-glutamate-specific TAXI SBP, BP0403.

**Figure 6.**
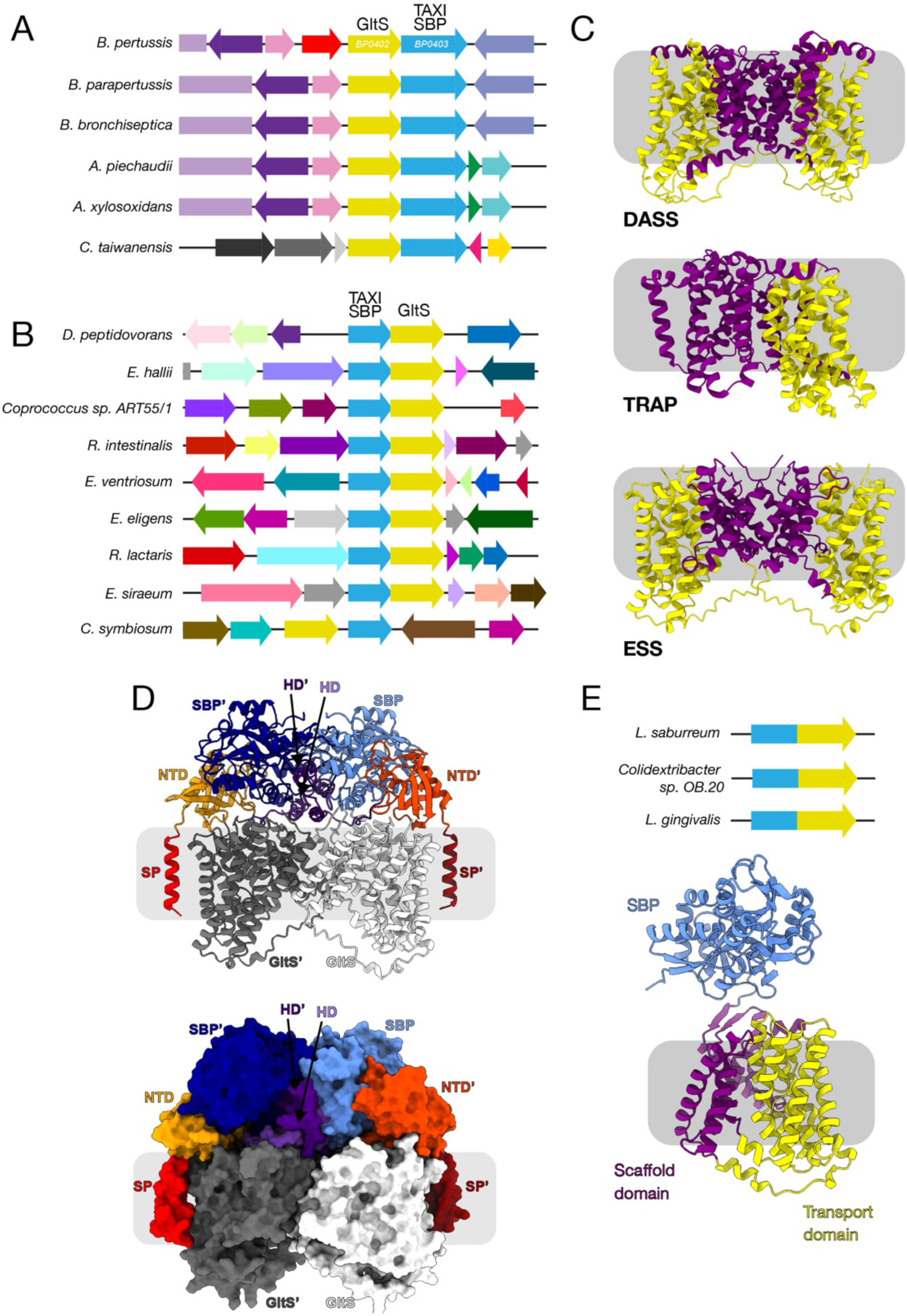
Linkage between genes encoding TAXI SBPs and GltS, and modelling of the TAXI-GltS interaction. **A)** Genome context of BP0403 and homologues (blue genes) from 5 other bacteria showing close proximity of gene encoding GltS (yellow gene). **B)** Genome context of genes encoding TAXI SBPs of conventional size (blue gene) and genes encoding GltS (yellow gene). C) Structural comparison of DASS transporter crystal structure (VcINDY, PDB: 5UL7), TRAP transporter cryo-EM structure (HiSiaQM, PDB: 7QE5), and ESS Colabfold model (BpGltS). Coloured to highlight scaffold (purple) and transport (yellow) domains. **D)** Cartoon representation (top) and surface representation (bottom) of the heterotetrameric BP0403:GltS complex modelled using AlphaFold Multimer. The domains of BP0403 are colour coded and labelled as in Fig 4. The GltS protomers are coloured white and dark grey. **E)** Examples of fusion between TAXI and *gltS* genes in the family Lachnospiraceae. Bottom: AlphaFold2 structural model of *Lachnoanerobaculum saburreum* fused TAXI-GltS. Coloured to highlight scaffold (purple), transport domain (yellow) and SBP (blue).

As there are no GltS structures available, we modelled the GltS dimer using Colabfold^27^, which revealed the presence of features strongly reminiscent of the Na^+^:succinate transporter VcINDY and TRAP transporter membrane components; namely, a scaffold domain and a transport domain characteristic of an elevator type mechanism (Fig. 6C)^16,17^. In addition, GltS has long been known to share a similar topology with the Na^+^:citrate transporter CitS, a member of the 2-hydroxycarboxylate (2HCT) family^36^, which operates via an elevator mechanism^37,38^. Based on the observation that both BP0403 and GltS are dimeric, we hypothesise that, upon binding glutamate, BP0403 presents the substrate to GltS, which uses an elevator mechanism similar to TRAP transporters, VcINDY and CitS to transport the substrate across the membrane.

To investigate the possible structural arrangement of the heterotetrameric BP0403:GltS complex, we modelled it using AlphaFold Multimer^28^. The combined pTM+ipTM score of about 0.6 indicates that we can reasonably predict the overall fold and protein/protein interactions of the complex. Interestingly, our BP0403:GltS model suggests that BP0403 undergoes a substantial domain rearrangement upon interaction with GltS compared to when the BP0403 dimer is modelled alone (Fig. 6D). For each BP0403 protomer in the tetrameric complex, the NTD interacts with the transport domain of one GltS protomer and the SBP domain interacts with *the other* GltS protomer. The second GltS protomer is arranged as a mirror image of the first protomer. The helical domains overlay the scaffold domain where they intertwine to produce a SBP domain swap. In the physiological setting, BP0403 would be attached to a lipid headgroup via an N-terminal cysteine residue, and our tetrameric model positions this cysteine residues precisely where we predict the lipid headgroups would arrange, supporting the feasibility of such an arrangement (Fig. 6D). Both SBP domains in our model are positioned above the scaffold:transport domain interface, which is the likely position of the GltS binding site. Therefore, our tetrameric BP0403:GltS model provides a mechanically and structurally feasible way for these proteins to work in concert during transport.

While the functional coupling between the TAXI SBPs and GltS needs to be experimentally verified, our hypothesis is substantially strengthened by the existence of several examples of *TAXI*-*gltS* gene fusions in genomes of the bacterial family Lachnospiraceae, which result in an N-terminal TAXI SBP domain fused to a C-terminal GltS (Fig. 6E). Presumably in this situation, GltS would still dimerise leading to a system reminiscent of the one we propose for BP0402/03.

## Discussion

In this work, we have described the discovery and characterisation of a novel type of glutamate-binding SBP in the TAXI family with a domain architecture completely distinct from all other TAXI proteins characterised to date. Moreover, it is genetically linked with, and translationally coupled to, the Na^+^:glutamate symporter GltS and not with a conventional TRAP transporter. Our results, combined with the observation that some bacteria possess single protein TAXI-GltS fusions, strongly suggest that this unprecedented arrangement represents an entirely novel type of binding-protein dependent secondary transporter.

BP0403 is much larger than conventional TAXI SBPs. We have shown by AlphaFold modelling and biochemical analysis that it is composed of separate and independently folded domains at the N- and C-termini, linked by a central helical domain. The distinctive double trough in the DSF profile of the full-length protein was suggestive of independent melting events of two domains with different Tm values, which we confirmed by analysis of the isolated N- and C-terminal domains. Evidence from both structural alignment of the C-terminal domain with the conventional-sized TAXI SBP VcGluP and its ability to bind substrate, unequivocally identifies this domain as the SBP element of the protein. Our modelling indicates that the helical domain forms the dimerization interface, which is supported by molecular weight analysis of the SBP and NTD in isolation (Fig. 5). An outstanding question is what function the NTD serves as this domain does not have sequence or structural similarity to any functionally characterised proteins.

In contrast to DctP-type SBPs, the substrates bound by TAXI proteins are far less well characterised. The first TAXI SBP structure to be determined had a bound ligand consistent with either glutamate or glutamine^21^, and subsequent characterisation of other TAXI SBPs demonstrate or suggest specificity for L-glutamate^22,39^, indicating this dicarboxylic amino-acid is a common ligand in this family. This is in keeping with our identification by DSF of L-glutamate as the ligand for BP0403. Moreover, we observed conservation of the key Y and Q residues in BP0403 that were shown to be crucial for L-glutamate binding in VcGluP^22^. However, characterisation of a TAXI-TRAP system from *Proteus mirabilis* revealed specificity for both 2-oxoglutarate and 2-hydroxyglutarate^23^, and a system from *Azoarcus* transports orthophthalic acid^40^, hinting at a greater diversity of substrates.

Most GltS proteins operate as Na^+^:glutamate symporters without the need for interaction with an SBP. BP0402 is the sole member of the ESS family present in *B. pertussis* but we do not yet know if it can operate independently of BP0403. What could be the physiological advantage of the interaction of these proteins? One possibility is that the SBP could bind a different ligand, thus effectively changing the substrate specificity of the membrane transporter. However, as we have shown that BP0403 binds L-glutamate and all characterised GltS systems transport glutamate, we find this unconvincing. A more likely explanation is that recruitment of the TAXI protein confers a higher affinity to the transporter compared with GltS alone. Indeed, our tyrosine fluorescence titrations of the isolated SBP indicate a sub-µM Kd for L-glutamate, whereas the Km of *E. coli* GltS for L-glutamate in membrane vesicles is 13.5-30 µM^35,41^.

A higher affinity transporter would be advantageous in the scavenging of scarce nutrients. In the tissues of mammalian hosts, extracellular L-glutamate concentrations are relatively low as this amino-acid is rapidly transported and metabolised by eukaryotic cells^42^. As *B. pertussis* adheres to and grows on cell surfaces, rather than being intracellular, the association of GltS with a TAXI protein to scavenge glutamate could result in a competitive advantage for this pathogen. Glutamate has in fact long been known to be of major importance in the physiology of *B. pertussis*. The bacterium is unable to utilise carbohydrates due to the lack of genes encoding key glycolysis enzymes and a small selection of amino-acids that are degraded to 2-oxoglutarate are the best carbon sources for growth^43^. Of these, glutamate enhances growth rate and cell yield the most and is the key component in the standard Stainer-Scholte cultivation medium^44,45^. Glutamate availability is known to modulate expression of many genes in *B. pertussis*^46^, and Glu limitation (as would be experienced in vivo) leads to up-regulation of some key virulence factors, such as the type 3 secretion system (T3SS) as well as enhanced auto-aggregation ability that may be important in adhesion^42^. Thus, understanding glutamate transport pathways in *B. pertussis* is key to fully appreciating the relationship between this amino-acid, colonisation and pathogenicity. In this regard, in addition to the TAXI-GltS system identified here, there is evidence that *B. pertussis* possesses several alternative glutamate transporters, including at least one ABC system with an SBP (BP3831) that is regulated by a small RNA^47^. The structures of two DctP-type SBPs (BP1887 and BP1891) of uncharacterised TRAP transporters have been determined, both of which bind pyroglutamate^48^. Pyroglutamate (5-oxo-L-proline) can be converted intracellularly to glutamate by pyroglutamase so this may represent an alternative strategy to obtain glutamate^49^. Finally, *B. pertussis* encodes a large number of genes encoding SBPs of the tripartite tricarboxylate transporter (TTT) family; referred to as “Bugs” (*Bordetella* uptake genes)^50^. Most of the corresponding SBPs are of unknown function and substrate specificity, but the structure of BugE was obtained with bound glutamate, implying that this might be part of a TTT-type glutamate uptake system^50^. All of these mechanisms for obtaining glutamate in *B. pertussis* employ SBPs, consistent with high-affinity uptake, although mutant studies will be needed to reveal which are of the greatest physiological relevance.

In a diverse range of bacteria, we found genes for both elongated and conventional sized TAXI SBPs closely linked to *gltS* (Fig. 6A and B). However, in some bacteria we identified TAXI-GltS gene fusions (Fig. 6E). Examination of the INTERPRO database reveals that of ∼10,000 entries for GltS, only 14 have this TAXI-GltS fusion architecture and the majority of these are found in members of one family, the Lachnospiraceae. These bacteria are poorly characterised members of the gut microbiome but they are firmicutes which therefore (presumably) do not have a conventional periplasm. Thus, the evolution of a TAXI-GltS fusion protein may be rationalised as one solution to ensure tethering of the SBP to the membrane transporter in these bacteria.

Prior to this study, the only families of secondary transporters known to operate with a soluble SBP were the TRAP and TT transporters. Why are SBPs not more widely associated with other diverse types of secondary transporters? It is now known that TRAP transporters operate by an elevator type mechanism. TT transporters are structurally very similar and likely operate in an analogous way. Our modelling suggests that GltS has clearly discernible scaffold and transport domains, implying that it too is likely to work as an elevator. This might suggest that some intrinsic feature(s) of the elevator mechanism is required to allow an associated SBP to bind and release its ligand as part of the transport cycle. The identification of further novel classes of SBP dependent secondary transporters will allow this proposal to be tested.

## Acknowledgements

DJK acknowledges the receipt of a Leverhulme Emeritus Fellowship (EM/2024-005) from the Leverhulme Trust, UK. This work was financially supported by a Biotechnology and Biological Sciences Research Council (BBSRC) grant (BB/V007424/1) awarded to CM and the SoCoBio BBSRC doctoral training partnership (BB/T008768/1). VL acknowledges that this research was completed in part with computational resources and technical support provided by the Research Computing Center at the Medical College of Wisconsin.

## Author contributions

LMJ purified proteins and collected binding assay and oligomeric state data. JFSD performed additional binding assays. CM, JFSD and DJK performed the bioinformatic analysis. VL modelled structural complexes and evaluated model quality and CM analysed the structural models. CM supervised the research. CM and DJK wrote the manuscript. All authors read and edited the manuscript.

## Data availability statement

All the data are contained in the manuscript.

## Supplementary Information

**Supplementary figure 1.**
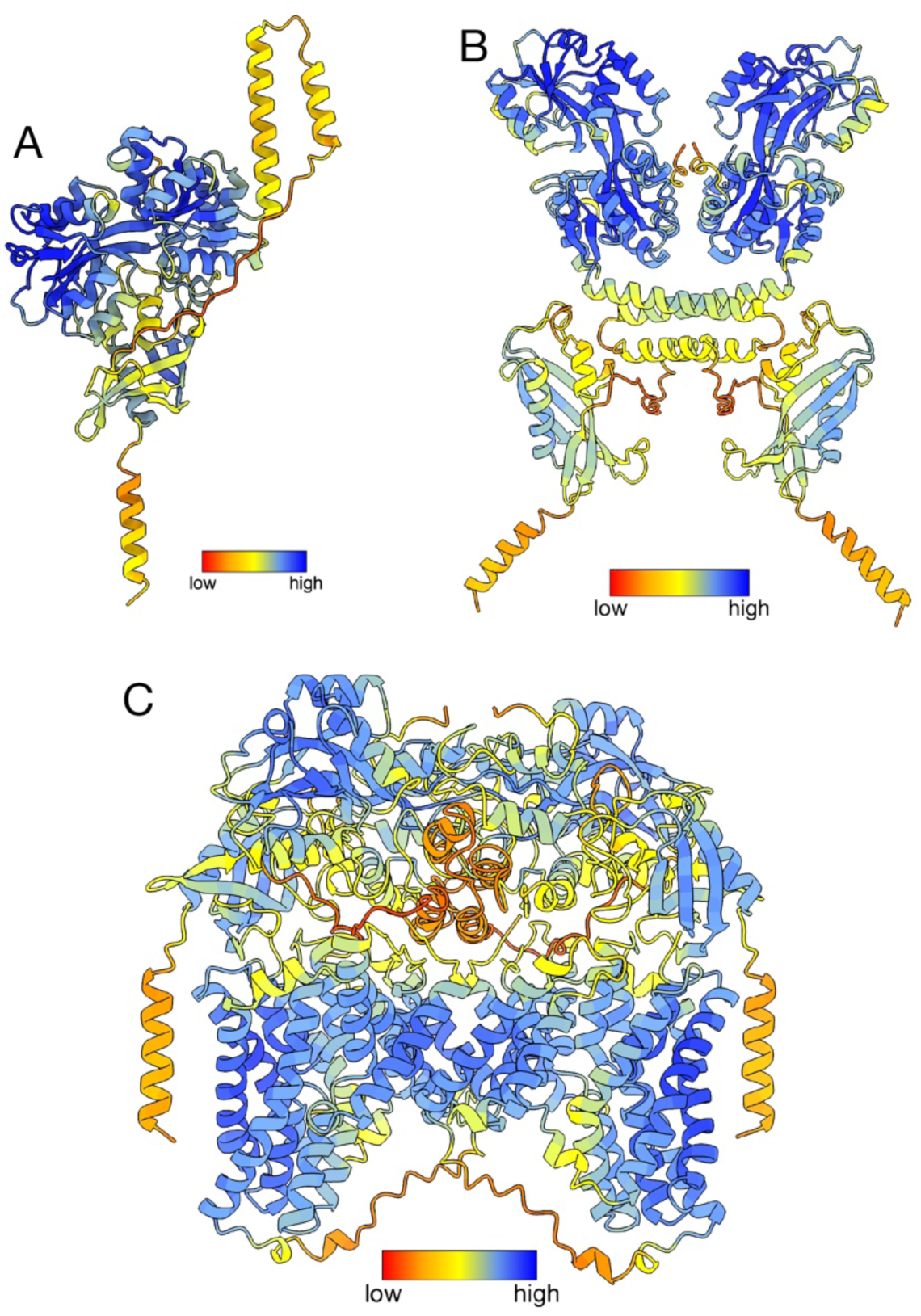
Predicted local distance difference test (pLDDT) scores. Models generated in study coloured by pLDDT score; **A)** BP0403 AlphaFold 2 model, **B)** BP0403 dimer ColabFold model, and **C)** BP0403:GltS heterotetramer AlphaFold Multimer model.

